# *pastclim*: an R package to easily access and use paleoclimatic reconstructions

**DOI:** 10.1101/2022.05.18.492456

**Authors:** Michela Leonardi, Emily Y. Hallett, Robert Beyer, Mario Krapp, Andrea Manica

## Abstract

The recent development of continuous paleoclimatic reconstructions covering hundreds of thousands of years paved the way to a large number of studies from disciplines ranging from paleoecology to linguistics, from archaeology to conservation and from population genetics to human evolution. Unfortunately, such climatic data can be challenging to extract and analyze for scholars unfamiliar with such specific climatic file formats.

Here we present *pastclim*, an R package facilitating the access and use of two sets of paleoclimatic reconstructions covering respectively the last 120,000 and 800,000 years. The package contains a set of functions allowing to quickly and easily recover the climate for the whole world or specific areas for time periods of interest, extract data from locations scattered in space and/or time, retrieve time series from individual sites, and easily manage the ice or land coverage.

The package can easily be adapted to paleoclimatic reconstructions different from the ones already included, offering a handy platform to include the climate of the past into existing analyses and pipelines.

## Background

Many research fields, such as ecology and archaeology, use climate reconstructions to contextualize information, such as the presence of a species at a given location in the past. Reconstructions for certain time points, such as the Last Glacial Maximum or Holocene, have been published and made publicly available for a number of climate models (*e*.*g*. Braconnot et al. 2012, Schmatz et al. 2015, Lima-Ribeiro et al. 2015, Kageyama et al. 2021). However, for certain analyses, time series are necessary, and over the last few years, a few datasets that provide climate reconstructions through time have become available (*e*.*g*. Armstrong et al. 2019, Brown et al. 2020, Beyer et al. 2020, Karger et al. 2021, Krapp et al. 2021, Timmermann et al. 2022).

The availability of such data paved the way for a large number of research questions among many different disciplines, *i*.*e*. ecology (Miller et al. 2021a, b), paleoecology (Leonardi et al. 2018, 2020, Somveille et al. 2020, Chen et al. 2021, Schap et al. 2021, Thorup et al. 2021), conservation (Beyer and Manica 2021), population genetics (Maisano Delser et al. 2021), archaeology (Racimo et al. 2020, Betti et al. 2020, Beyer et al. 2021, Krzyzanska et al. 2021, Park and Marwick 2022, Cerasoni et al. 2022, Timbrell et al. 2022), the evolution of the genus *Homo* (Will et al. 2021, Timmermann et al. 2022), anthropology (Leonardi et al. 2017, Padilla-Iglesias et al. 2021) and linguistics (Beyer et al. 2019).

A challenge for the use of such climatic datasets covering time series is that they are large in size, requiring the use of specialized software to manipulate and extract the data. Climate reconstructions are generally stored in the netCDF format, and the *cdo* (Schulzweida 2021) tools in bash, and their implementations in R and Python can be used to directly access the data. Software designed to handle raster data, such as the R packages *raster* (Hijmans 2022a), and *terra* (Hijmans 2022b), can directly extract data from netCDF files, but dealing with files with hundreds of time steps and dozens of variables can be challenging. Furthermore, different authors have packaged their data in different ways, either as a single large netCDF file, or broken down into multiple files that have to be downloaded from various public repositories. The naming of variables is also often inconsistent, adding a further challenge for users.

Here we present *pastclim*, an R package specifically designed to download and manipulate two such datasets: one from Beyer et al. 2020, which covers the last 120,000 years at intervals of 1,000-2,000 years based on Global Circulation Models HadCM3 (Singarayer and Valdes 2010) and HadAM3H (Valdes et al. 2017), and one from Krapp et al. 2021, which covers the last 800,000 years at intervals of 1,000 years based on a statistical approach to reconstruct HadCM3 outputs. For both datasets, we provide 22 bioclimatic and vegetation variables, bias-corrected and downscaled at a spatial resolution of 0.5° × 0.5°. We note that the original datasets also include monthly temperature and precipitation estimates, but these are not currently handled by *pastclim*. Furthermore, the datasets were trimmed to mask the ice sheets, as reconstructions under ice caps are not useful for most ecological and anthropological analyses and masking such large datasets can be time-consuming.

The functions of the package are generic and can be used to process similar datasets.

### Description of the data

*pastclim* allows direct access to two datasets: “Beyer2020” covering the last 120,000 years; “Krapp2021” which goes back to 800,000 years ago. These datasets must be downloaded using functions from the package. Additionally, a small “Example” dataset, which includes a few time points from Beyer2020, is included in the R package for use in the vignette and testing.

The two datasets include all WordClim variables (Hijmans et al. 2005) except bio02 (Mean Diurnal Range) and bio03 (Isothermality), and also contain annual net primary productivity and biome reconstructions. The latter is a categorical variable representing the type of natural vegetation, reconstructed by the BIOME4 model (Kaplan et al. 2003). In addition to the bioclimatic variables presented in the original publications (Beyer et al. 2020, Krapp et al. 2021), the updated datasets now also include two measures of topography, namely altitude and rugosity, based on the ETOPO1 dataset (NOAA National Geophysical Data Center 2009). The latter was calculated as the standard deviation of the altitude of all above-water 1’ x 1’ cells included in each of the 0.5° × 0.5° cells as used in the climate reconstructions.

The original Beyer2020 and Krapp2021 datasets are freely available to download from Figshare (https://figshare.com/articles/dataset/LateQuaternary_Environment_nc/12293345/4) and OSF (https://osf.io/8n43x/). They are stored in two different ways, as a large netcdf file containing most variables for Beyer2020 (plus an additional small file for some additional vegetation variables) and as separate netcdf files for each variable for Krapp2021. The repackaged data for the two datasets are found on figshare (https://figshare.com/articles/dataset/pastclim_beyer2020_v1_0_0/19723405 and https://figshare.com/articles/dataset/pastclim_krapp2021_v1_0_0/19733680)

## Methods and features

The *pastclim* package allows users to download and manage the datasets using a unified interface. The user can either store the datasets in a custom directory, or use a companion package, *pastclimData*, to store the data within the package itself and find them automatically without the need of setting a path.

We envision two main uses for *pastclim*. One on hand, a user might have one (or a few locations) of interest for which they want to retrieve a time series of bioclimatic variables. Under this scenario, such time series are simply retrieved through the function *time_series_for_locations()*, and they can then be compared to other information for those sites. In **Applied Example 1** we provide a step-by-step illustration of how (zoo)archaeologists might use *pastclim* to provide context for their habitat reconstructions based on faunal data from different stratigraphic layers of a site.

The other use we envisage is for larger-scale analyses, where a user might be interested in climatic reconstructions for a large number of sites, likely sampled at different times, and possibly contrast this information with the background climate across a whole region. In *pastclim*, the functions *climate_for_locations ()* and *climate_for_time_slice()* provide these functionalities. **Applied Example 2** provides an example of how these functions can be combined to explore the bioclimatic niche of a species.

We also include several helper functions that facilitate the manipulation and visualisation of data. Data can be easily cropped to preset geographic extents that represent specific continental masses. Furthermore, the climatic fluctuations observed in the past not only led to changes in the bioclimatic variables but also significantly affected the extent of the permanent ice sheets and the sea levels, hence modifying the coastlines. In *pastclim* it is possible to get the ice and land masks for a given time slice using the functions *get_ice_mask()* and *get_land_mask()*. As mentioned earlier, note that the bioclimatic variables are not provided for cells that were under either water or ice sheets.

When it is useful to subdivide the data into Marine Isotope Stages (MIS), *pastclim* offers the function *get_mis_time_steps()* can get a list of time steps available for a given MIS, based on the subdivision proposed by Lisiecki and Raymo 2005. This, in turn, can be used to split the climatic reconstructions into MIS to perform some basic plotting or further analyses.

**Table 1.** Main functions of pastclim

| Name | function |
| --- | --- |
| *get_vars_for_dataset()* | Download variables |
| *get_downloaded_datasets()* | Summary of the downloaded variables |
| *climate_for_locations()* | Get the climate for given locations (by coordinates) |
| *time_series_for_locations()* | Get a time series of the climate for given locations (by coordinates) |
| *climate_for_time_slice()* | Get the climate worldwide for a given time step |
| *get_ice_mask()* | Get the ice mask for a given time step |
| *get_land_mask()* | Get the land mask for a given time step |
| *get_mis_time_steps()* | Get a list of time steps available for a given MIS |

**Table 2.** Characteristics of the two databases accessible from pastclim

|  |  |  |
| --- | --- | --- |
| Wordclim variables | bio01: Annual mean temperature<br>bio04: Temperature seasonality<br>bio05: Minimum annual temperature<br>bio06: Maximum annual temperature<br>bio07: Temperature annual range<br>bio08: Mean temperature of the wettest quarter<br>bio09: Mean temperature of driest quarter<br>bio10: Mean temperature of warmest quarter<br>bio11: Mean temperature of coldest quarter<br>bio12: Annual precipitation<br>bio13: Precipitation of wettest month<br>bio14: Precipitation of driest month<br>bio15: Precipitation seasonality<br>bio16: Precipitation of wettest quarter<br>bio17: Precipitation of driest quarter<br>bio18: Precipitation of warmest quarter<br>bio19: Precipitation of coldest quarter |  |
| Vegetation variables | Net primary productivity<br>Leaf area index (Beyer2020 only)<br>Biome |  |
| Topography variables | Altitude<br>Rugosity |  |
| Time slices | Beyer2020: Every 1,000 years between the present and 22,000 BP, every 2,000 years between 22,000 BP and 120,000 BP | Krapp2021: Every 1,000 years between the present and 800,000 BP |

### Examples

#### Applied example 1

The first example shows how to use *pastclim* for climate reconstructions for a single archaeological site with dated stratigraphic layers, *i*.*e*., in one location for multiple time slices. Example 1 shows how habitat reconstructions from vertebrate faunal data compare with climate reconstructions from *pastclim*. Chronometric dating data for the archaeological deposits containing vertebrate faunal remains from Contrebandiers Cave, Morocco, was extracted from Dibble et al. 2012. Each identified species of Artiodactyla from Contrebandiers Cave was assigned a habitat and feeding preference. The relative abundance of forest, savanna, and mixed forest/savanna Artiodactyla species was calculated for each archaeological layer following the number of identified species presented in Hallett 2018 (figure 1).

**Figure 1:**
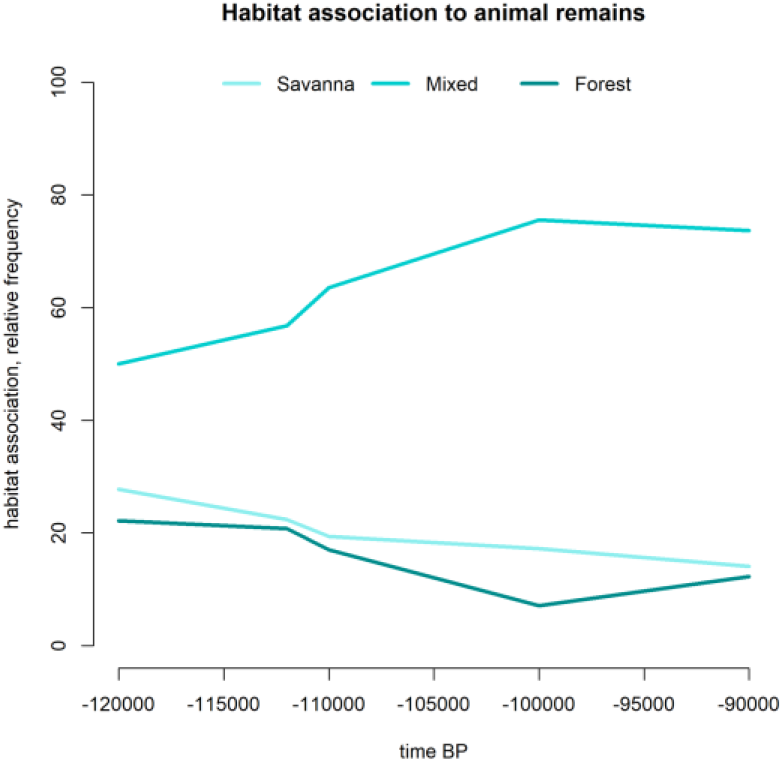
Relative abundance of habitat associated with Artiodactyla species from Contrabandiers Cave, Morocco, between 120,000 and 90,000 years ago.

The input (Supplementary material) is a table containing time (*i*.*e*. the chronometric age of the stratigraphic layer) and the relative percentage frequency of savanna, mixed, and forest Artiodactyla species. Climate can then be extracted for each layer with faunal data using the decimal coordinates of Contrebandiers Cave. As shown in figure 2, annual precipitation more closely follows the trajectory of change in Artiodactyla habitat types than the mean temperature of the warmest quarter. In this example, vertebrate faunal data as a proxy for past habitat reconstructions agrees well with the annual precipitation climate variable extracted from *pastclim*.

**Figure 2:**
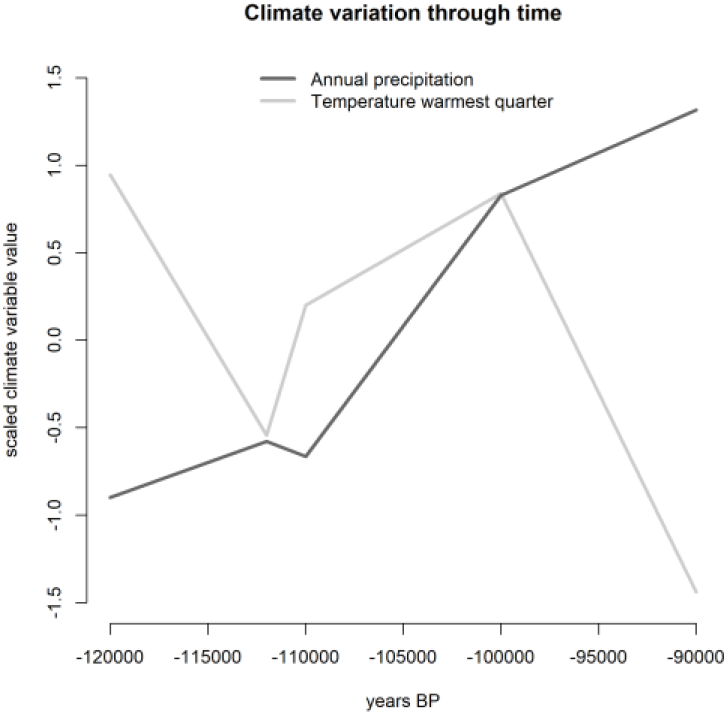
Variation of climatic variables through time in the region of Contrabandiers Cave, Morocco, between 120,000 and 90,000 years ago.

#### Applied example 2

The second example shows how to use *pastclim* to analyze a larger number of sites with lower individual coverage, *i*.*e*., a number of points scattered in space and time. The analyses presented will focus on comparing the climate associated with observations to the one from the whole region of interest, and they may represent exploratory steps prior to species distribution modelling (e.g. Miller et al. 2021b).

The input (Supplementary Material) is a simple table containing calibrated radiocarbon dates associated with horse remains from two different time steps (10,000 and 20,000 years ago), together with the associated geographic coordinates and site names. The data is extracted from (Leonardi et al. 2020) (figure 3) and the analyses are performed using the “Example” climatic dataset.

**Figure 3:**
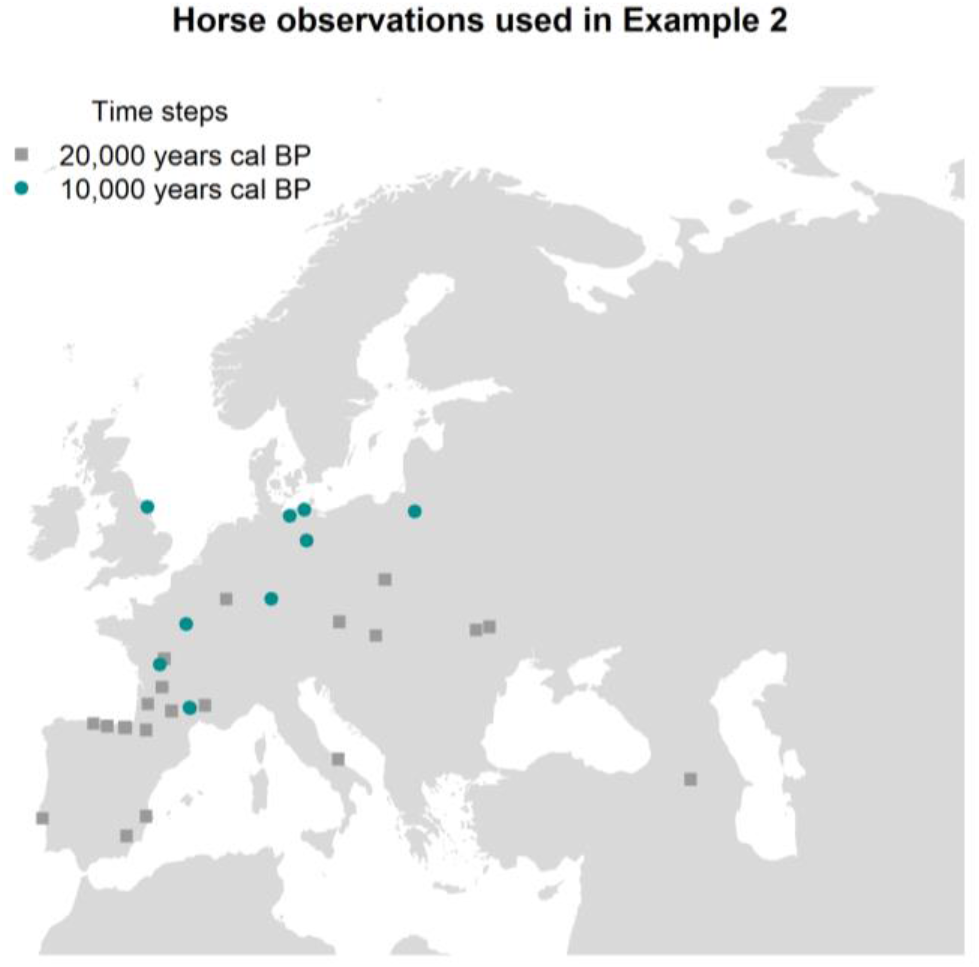
Map of the horse observations used for the analyses presented in Example 2

After transforming the calibrated radiocarbon dates into the associated time steps, it is straightforward to download the bioclimatic variables associated with the observations using *climate_for_locations ()*. It is then necessary to retrieve climatic information for the background, which can be done with the *climate_for_time_slice()* function. As the observations only cover Europe, it will then be necessary to crop the resulting climatic reconstruction to the right extent, that is already available within *pastclim* together with other macro-areas worldwide (*e*.*g*. Eurasia, Africa, Oceania, etc.).

Such outputs are given as tables and hence can be easily manipulated to perform standard analyses. For example, it is possible to compare in a lot of different ways the climate associated with horse occurrences and the whole of Europe. In the attached code (supplementary material) we show how to produce a PCA (figure 4a) and a plot comparing the density for each variable in Europe and in the horse data (figure 4b). This can be very useful, as the ones for which the distribution in the species differs from the background are likely to play a bigger role in shaping the distribution (Miller et al. 2021a).

**Figure 4:**
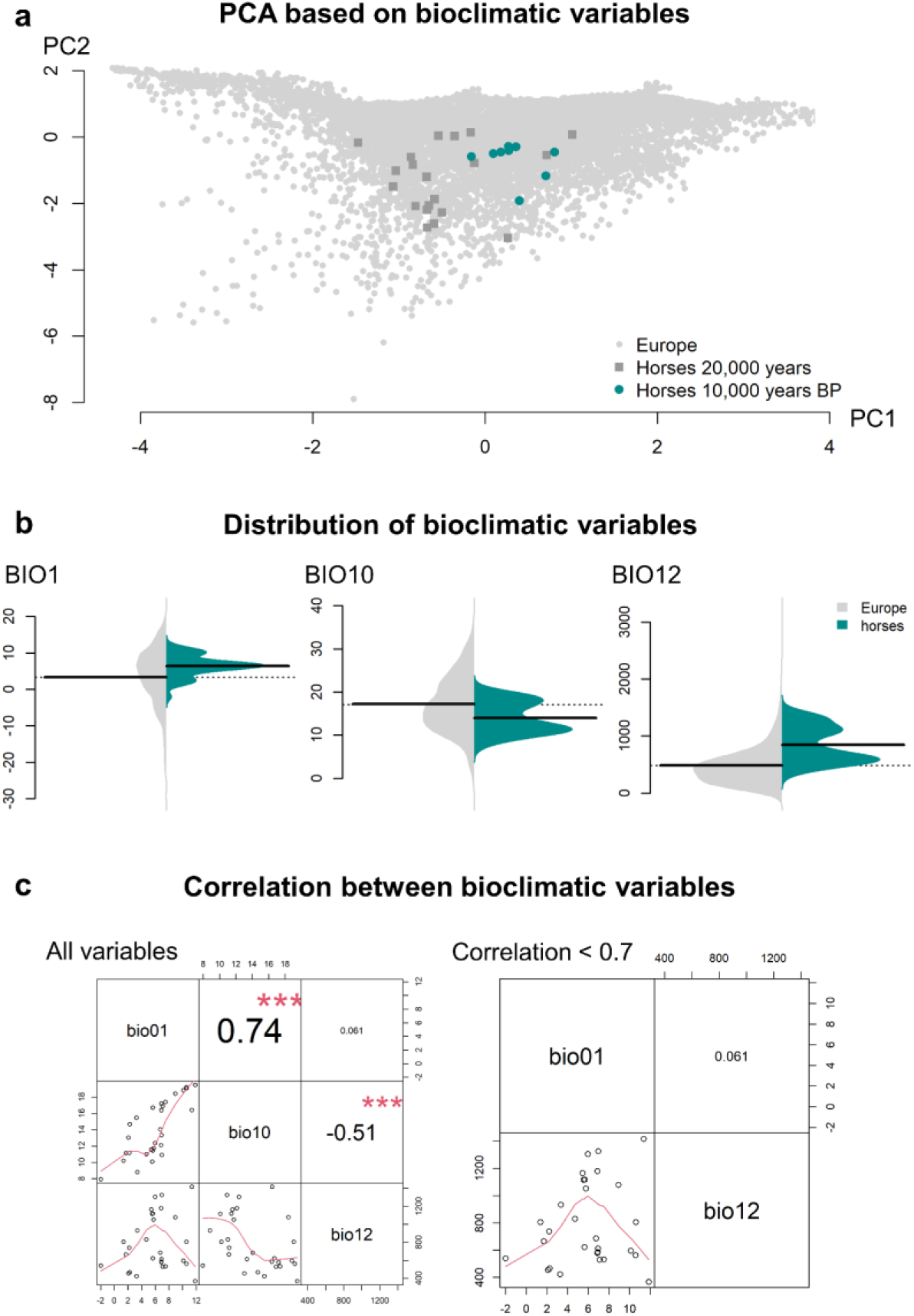
Analyses performed in Example 2. a: Principal Component Analysis based on climatic variables. b. distribution of climatic variables in locations occupied by horses (right, in blue) and in the whole of Europe (left, light grey) during both time slices analyzed. c: correlation between climatic variables in locations where horses have been observed in both time slices analyzed.

For species-distribution-model-like analyses (Elith and Leathwick 2009) it may be necessary to remove climatic variables showing a correlation in the data above a certain threshold (e.g. 0.7, Guisan et al. 2017). Given the standard format of the outputs from *pastclim*, it is straightforward to calculate the cross-correlation between the climatic variables in the observations and remove highly correlated ones using functions from existing libraries (figure 4c).

## Discussion

The *pastclim* package offers scholars studying the distant past (*e*.*g*., archaeologists, archeozoologists, paleoecologists) easily accessible climate variables from the past. Past changes in climate are often presented as the drivers of technological innovation and cultural change within archaeology (Richerson et al. 2009, Taller and Conard 2022). However, climate reconstructions of the past are often based on proxy data from locations distant from an archaeological site (as discussed in Jacobs et al. 2012, Burke et al. 2021). Local, site-specific climate reconstructions give archaeologists the opportunity to see how changes in climate and/or biome align with changes in human behavior for a given location.

Past climatic fluctuations heavily influenced the distribution and demography of humans and animals (Leonardi et al. 2018, Mondanaro et al. 2020, Fordham et al. 2022, Timmermann et al. 2022, Rodríguez et al. 2022). This is why *pastclim* can be a useful resource for scholars working in the field of (archeo)genomics, as it allows to easily extract climatic data to be integrated with genetic and genomic evidence when reconstructing the past (Lorenzen et al. 2011, Leonardi et al. 2017, Racimo et al. 2020, Theodoridis et al. 2020, Miller et al. 2021a, Maisano Delser et al. 2021).

Finally, the current rate of climatic changes due to human activity (Quintero and Wiens 2013) brings up the urgent need to understand how species react to climatic changes. Paleoclimate can be used as a virtual lab to (1) assess the extent to which species are able to change their ecological niche in response to environmental turnovers (Leonardi et al. 2020); (2) estimate the existing niche (Jackson and Overpeck 2000) though time, which can help to identify potentially suitable areas not apparent from the current distribution (3); identify Anthropocene refugia *i*.*e*. areas that provide protection from human activities and that will remain suitable in the long-term (Monsarrat et al. 2019); (4) more in general, gain a better understanding of the biological processes that allow surviving abrupt environmental changes (Fordham et al. 2020) which in turn may help to define better conservation strategies for the future.

### Package installation and availability

This package and the associated vignette and examples are free and open-source and are available for download on the GitHub page of the Evolutionary Ecology Group https://github.com/EvolEcolGroup/pastclim, where can also be found details about the license and the dependencies.

To cite *pastclim* or acknowledge its use, please cite the present work.

## Supplementary material

The input data and commented code needed to run the applied examples can be downloaded from https://doi.org/10.6084/m9.figshare.19790335

## Acknowledgements

We thank Megan McCrory, Johanna Paijmans, Sidney Leedham and Cecilia Padilla-Iglesias for testing the package during its development.

